# RichPathR: a gene set enrichment analysis and visualization tool

**DOI:** 10.1101/2023.08.28.555198

**Authors:** Binod Regmi, Stephen R. Brooks, Andrew J. Uhlman, Hong-Wei Sun

## Abstract

Gene set enrichment analysis (GSEA) is an important step for disease and drug discovery. Genomic, transcriptomics, proteomics and epigenetic analysis of tissue or cells generates gene lists that need to be further investigated in the known biological context. The advent of high-throughput technologies generates the vast number of gene lists that are up or down regulated together. One way of getting meaningful insights of the relationship of these genes is utilizing existing knowledge bases linking them with biological functions or phenotypes. Multiple public databases with annotated gene sets are available for GSEA, and *enrichR* is the most popular web application still requiring custom tools for large-scale mining. *richPathR* package is a collection of R functions that helps researchers carry out exploratory analysis and visualization of gene set enrichment using *EnrichR*.

**Availability:** The package, test data and additional figures can be downloaded from https://github.com/niams-bdmds/richPathR.git.

## 1. Introduction

High throughput sequencing and analysis of transcriptomic and epigenetic data sets often result in multiple gene sets associated with corresponding phenotypes of interest (Fortunel et al., 2003, Mootha et al., 2003, Subramanian et al., 2005). Further investigation is required to put these associations in biological context. There are several tools, annotated databases and web applications developed for similar analysis (Subramanian et al, 2007, Kuleshov et al., 2016). EnrichrR (Kuleshov et al., 2016) is one of the most popular web applications for annotating gene sets with biological function. EnrichR provides several mining, sorting and visualization options for querying a limited number of gene sets and libraries. This web based tool is convenient for analyzing small numbers of gene lists but less so for large scale mining and custom visualization of multiple gene sets and databases.

This R package, richPathR, has been developed as an extension of Enrichr web application for rapid mining multiple gene sets and libraries and exploratory visualization. Users can extract a large data frame as well as tables of the most ubiquitous and unique terms or pathways and customize their exploratory visualization.

## 2. Methods

This package is written to provide an easy-to-use tool for the users with only a basic computational background. The package provides clear guidance for navigating through building a large data frame to customized visualizations.

### 2.1 Test data

We used four gene sets obtained from expression profiling in tumor-associated endothelial cell of invasive ovarian cancer (Lu et al., 2007) and four libraries Cancer_Cell_Line_Encyclopedia, NCI-60_Cancer_Cell_Lines, NCI-Nature_2016, UK_Biobank_GWAS_v1, and KEGG_2021_Human for testing the script.

### 2.2 Calling the packages and dependencies

The users can load all the dependencies and packages using call_all_packages() function. RichPathR calls the enrichR package, an R package which provides programmatic access to EnrichR website functionality.

### 2.3 Generating a large data frame from multiple gene sets and database libraries

The package implements enrichr package and generates several excel files which contain one sheet with nine columns for each database library. Out of these nine columns, five columns are used for generating the data frame. Generating these data frames compiling all terms across all gene sets and libraries is the most useful functionality of this package. Researchers can extract very useful information from these data frames quickly. Researchers can use functions within this package or their own script for analysis and visualization of this data frame.

Two data frames are generated for exploratory visualization. The smaller data frame created by enrichr_df() has a single row for each term. The larger data frame created by expanded_enrichr_df() explodes the gene set for each term separating them into each row.

The expanded data frame is used for exploratory visualization like enrichr_heat_map(), gene network, and tile plot where researchers are interested in the top hit to individual gene level details.

### 2.4 Generating tables of unique and ubiquitous terms/pathways

Mining multiple database libraries for multiple input gene sets, the package generates two useful tables of the most ubiquitous terms (max(count)) and unique (count = 1) terms. The first table is usually smaller than the second. The table of unique terms helps to explore the rare hits or recently added terms in the database. The ubiquitous table shows the most common pathways that have been recorded in multiple gene sets and libraries.

### 2.5 Estimating Jaccard’s index

The function estimate_jaccard_similarity() adds one column to existing data frame. Jaccard’s index is calculated by the ratio of intersection and union of input gene set and gene set of the data base. This calculation is useful for quickly sorting the terms with higher overlapping gene sets.

### 2.6 Estimating adjacency matrix

The adjacency matrix is estimated to store node and edge information to visualize term’s(pathways or phenotypes) interaction with multiple cluster of genes. The edge list is extracted from extended data frame and undirectional adjacency matrix is calculated. The matrix is visualized using igraph function graph_from_adjacency matrix.

### 2.7 Visualization

#### 2.7.1 Terms/pathways related visualization

The distribution of terms/counts across the libraries is visualized using single and mixed bar plots after filtering the most significant terms/pathways based on combined score (Figure 1). The enrichr database provides three ranking methods for terms:. Fisher’s exact test (FET) p-value, rank deviation z-score and combined score. The combined score is the log product of p-value and z-score. FET p-value is biased on length of the input gene set whereas z-score alone is slightly better than combined score, however we selected combined score for sorting the terms and downstream analysis due to its mathematical computability. Users can select their minimum combined score; the default is 5.

**Figure 1:**
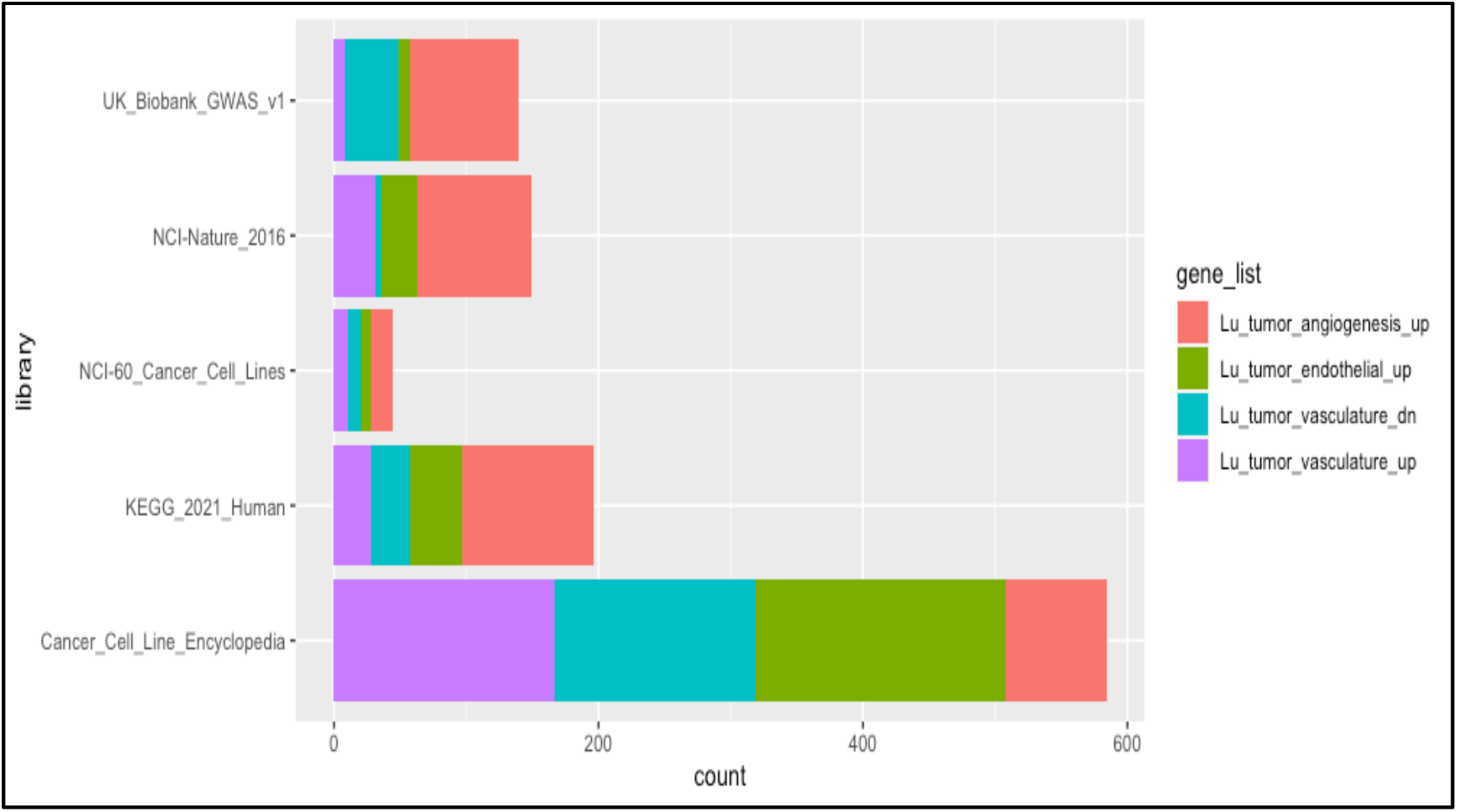
Pathways/terms presented in the gene sets and libraries.

Unique or ubiquitous distribution of terms across gene is visualized using horizontal bar plot. The violin plot is particularly useful to extract and visualize the most significant terms based on combined score (Figure 2, Figure 3). Figures 2 and 3 visualize where the combined scores overshoot. The first and third gene set show more significant terms as compared to second and fourth gene set (Figure 2). Between these two gene sets, the first gene set is shared in KEGG library whereas both first and third gene sets are shared in UK Biobank library (Figure 3). Our test data is expression profiling of tumor-associated endothelial cells of ovarian carcinoma, so the top hit term based on combined score is ‘pathways in cancer’ with 16 genes of input gene set overlapped with library gene set of 531 genes. This pathway is recorded in KEGG_2021_Human library. This exemplifies how quickly we can reach to potential terms using this package for further investigation. The smaller data frame (enrichr_df()) is used for all these visualizations.

**Figure 2:**
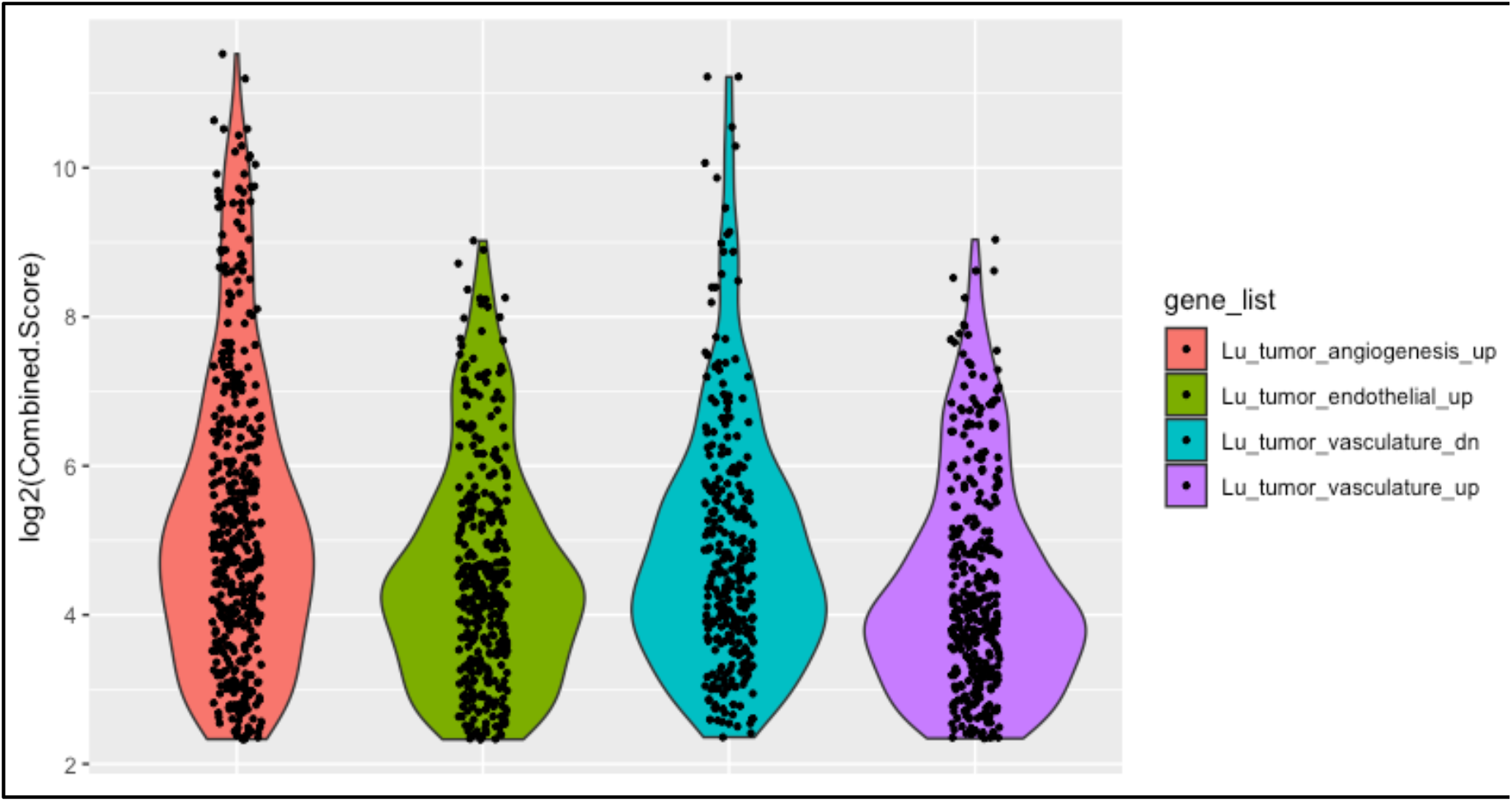
Signifincat pahways/terms extracted for the gene lists.

**Figure 3:**
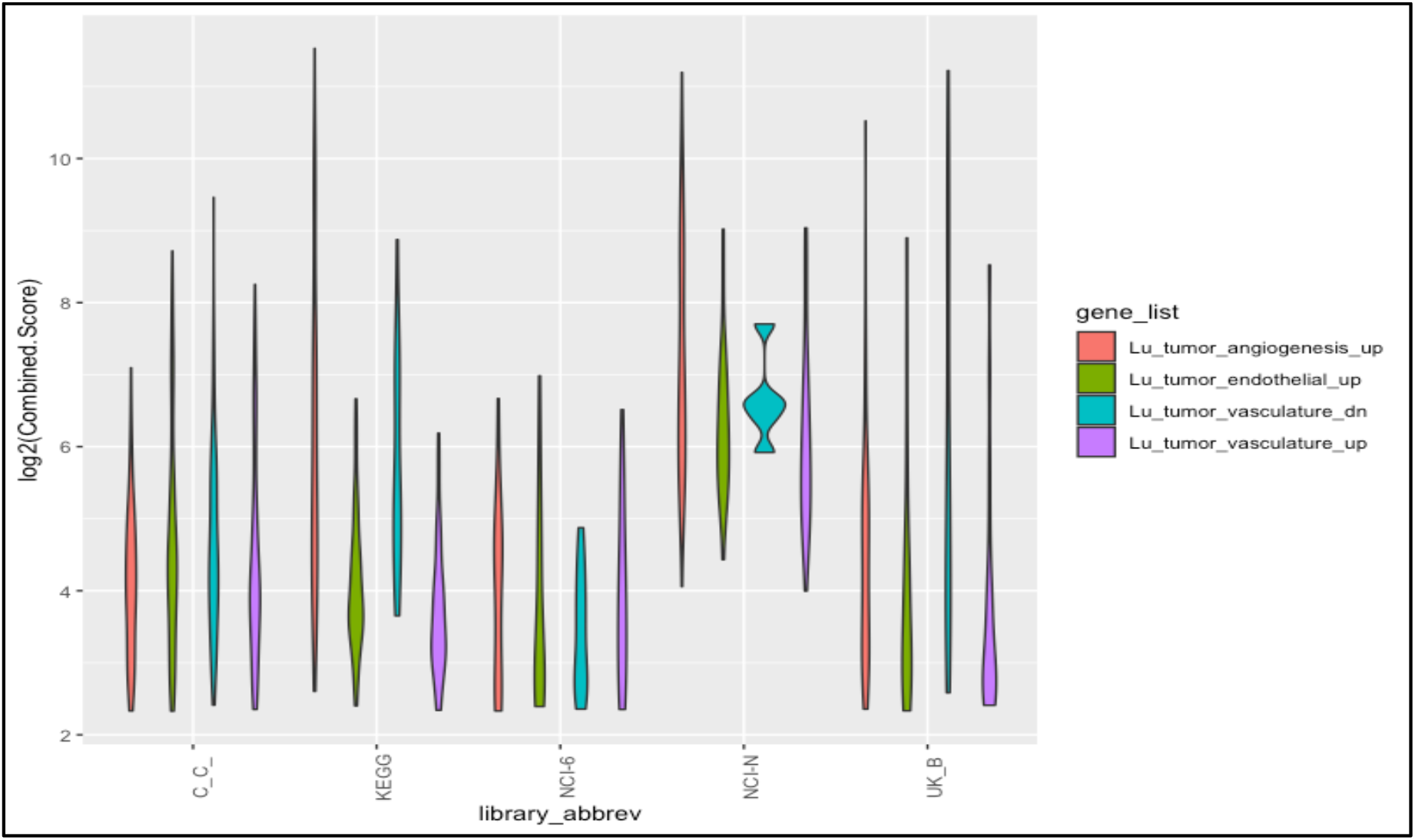
Significant pathways/terms extracted for the gene lists and libraries.

#### 2.7.2 Gene level visualization

The enrichr_heat_map() and tile_plot() functions are used for visualization at an individual gene level. The enrichr_heat_map visualizes the top 30 high frequency genes distributed across gene sets and libraries (Figure 4). The tile plot visualizes a heat map of count of terms or pathways in input gene sets and libraries. Here, the heat map (Figure 4) shows CCND2 is the hub gene that produces cyclin D2 protein. Cyclin D2 controls rate of cell growth and proliferation in the body.

**Figure 4:**
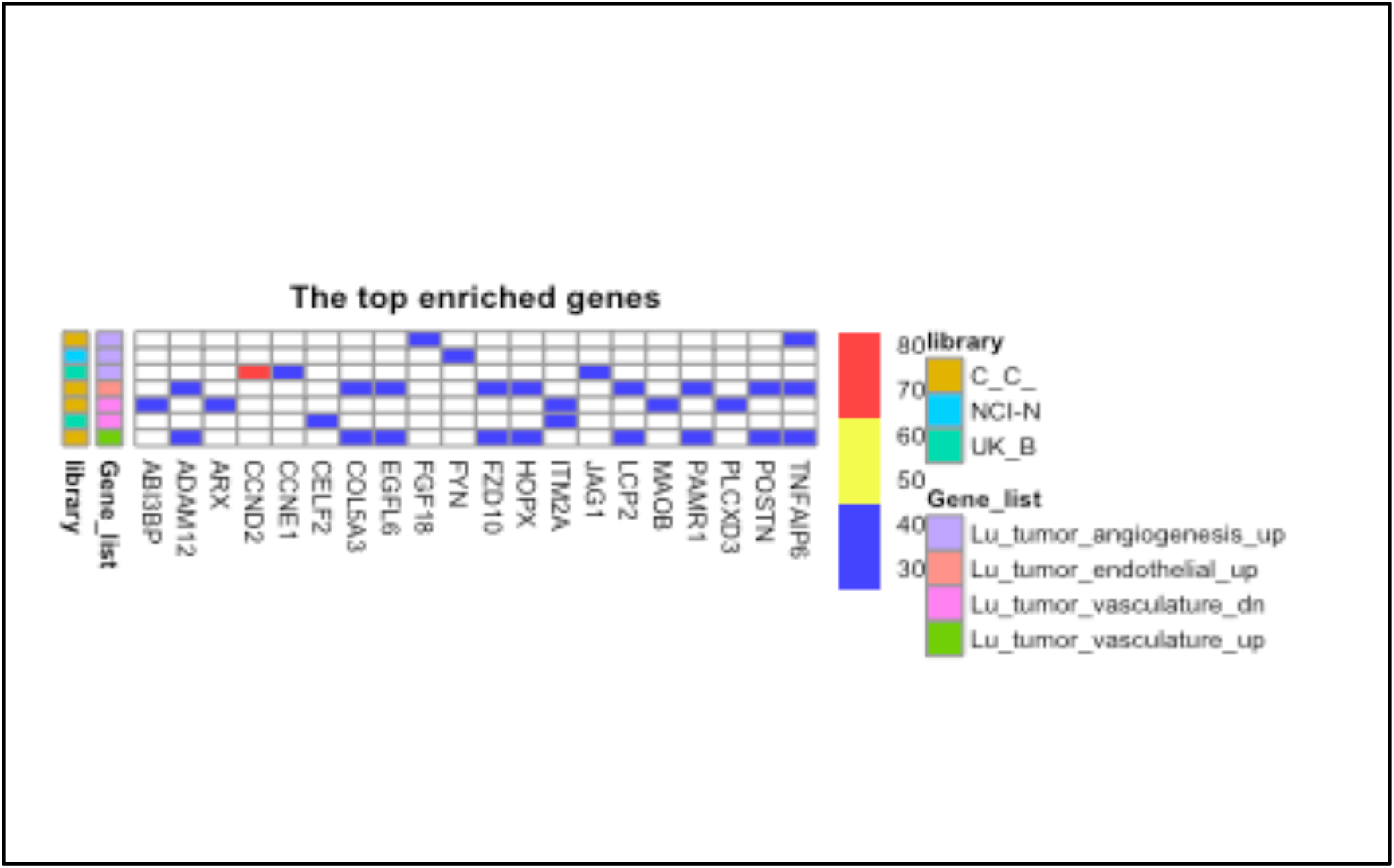
Hub (high frequency) genes presented in one or multiple gene sets and libraries.

#### 2.7.3 Visualizing terms(pathways) and gene network

For visualizing the interaction of terms and multiple gene sets, network_plot can be used. It requires a list of terms and the expanded data frame (Figure 5). We selected three pathways: FoX, Wnt, and Toll-like receptor from our test data. The network plot shows that CCND2 gene is presented in both Wnt and FoxO signaling pathways where as MAPK3 and PIK3CB are presented in FoxO and Toll-like receptor pathways. The network plot is useful to visualize the interaction of gene sets and particular terms presented in the specific database.

**Figure 5:**
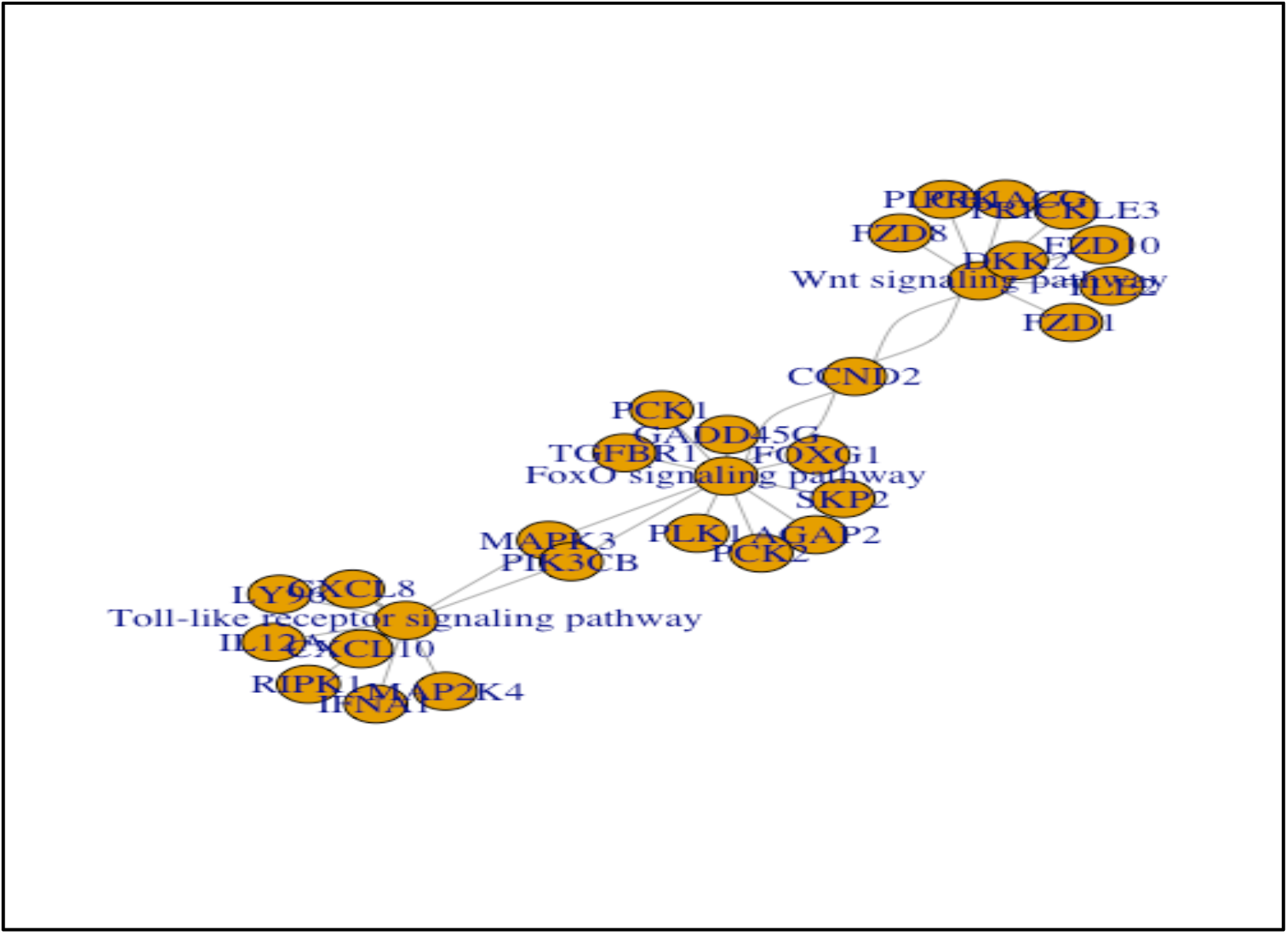
Interaction network among terms and genes for the selected terms.

## 3 Discussion

The polygenic mode of genotype-phenotype relationships has been the subject of discussion in the current trend of growing high throughput genetic, genomic, transcriptomic and epigenetic data (Berg and Coop, 2014; Chesler, et al., 2014). Analyzing the associated gene sets and creating annotated databases is ever growing (Liberzon et al., 2015; Subramanian et al., 2007; Yoo et al., 2015; Wang et al., 2013). There are multiple web applications that provide software tools and databases to conduct gene set enrichment analysis (Glaab et al., 2012, Wang et al., Merico et al. 2010).

The current package fills the gap of available tools for rapid mining of pre-annotated data of pathways/terms annotated from different high throughput experiments (bulk RNA/ ATAC/ChIP-seq). A single transcriptomic or epigenetic high throughput sequencing experiment might generate several gene sets andmining these gene sets one at a time could be time consuming. The current package provides rapid mining and exploratory visualization tools. The single data frame compiling all gene sets and annotated terms is very useful functionality for rapid data mining. The table of ubiquitous terms helps to extract the gene sets supported by multiple data sources. Using the violin plot for example, we can rapidly scan the most significant terms or pathways distributed in input gene sets and data base libraries. The heat map could be useful for uncovering hub genes in similar gene sets in related libraries. The network plot provides a useful tool for mining and visualizing highly enriched gene or a set of genes. This plot could also be useful to understand and visualize interaction of interesting terms (i.e. pathways, cell lines or phenotypes) presented in the database. The exploratory analysis of available libraries for large input gene sets points correct direction for further studies, provides ground for generating testable hypotheses and saves investigator’s time.

## Acknowledgement

This research was supported by the Intramural Research Program of the National Institute of Arthritis and Musculoskeletal and Skin Disease(NIAMS) of the National Institutes of Health(NIH). This work utilized the computional resources of the NIH HPC Biowulf cluster.

